# Neuromark Fusion: A Replicable Multimodal Template for Structure-Function Fusion of Brain MRI

**DOI:** 10.64898/2026.01.23.701328

**Authors:** Marlena Duda, Bradley Baker, Jessica Turner, Theo Van Erp, Vince Calhoun

## Abstract

Multimodal data fusion is a powerful technique for extracting shared and complementary information about the brain that is captured across neuroimaging modalities. Independent component analysis (ICA)-based approaches are among the most widely utilized methods for multimodal fusion, as they are data-driven, robust to noise, and capable of identifying complex, hidden linkages of varying strengths across high-dimensional datasets. However, the data-driven nature of ICA fusion approaches can make comparisons across analyses difficult without a normative framework in place. In this work, we utilize resting state functional MRI (rsfMRI) and structural MRI (sMRI) scans from >15,000 subjects to generate a normative model of multimodal structure-function linkages that can be used as a template to guide ICA fusions of new datasets. When applying this template in two datasets, resultant components exhibit high correspondence to the template even in small sample sizes, and subject-level loadings from template-derived ICs show significant associations to age.

## 1. INTRODUCTION

Multimodal fusion has emerged as a powerful framework for characterizing normative patterns of brain organization that span structure and function. By integrating complementary information from modalities such as structural MRI (sMRI) and functional MRI (fMRI), multimodal approaches enable the estimation of latent axes of population-level variation that are not observable within any single modality. ICA-based fusion methods, including joint ICA [1], parallel ICA [2], mCCA + jICA [3], and linked ICA [4], provide data-driven means of modeling shared sources of inter-subject variability across high-dimensional neuroimaging data. When applied at scale, these methods can be interpreted as normative models that capture typical covariation patterns in brain structure and function, against which individual deviations may be quantified. Such normative representations are increasingly central to efforts to map brain-behavior relationships, identify clinically relevant deviations, and support precision neuroscience.

A key limitation of fully data-driven fusion approaches, however, is that the resulting independent components (ICs) may lack reproducibility and comparability across datasets, hindering their use as stable normative references. Constrained ICA methods address this challenge by incorporating spatial priors that enforce correspondence across analyses, yielding a fixed set of ICs that can serve as a template. While normative templates have been developed and validated for unimodal fMRI data [5–8], an analogous multimodal normative model has not yet been established. Here, we leverage resting state fMRI and sMRI data from over 15,000 participants in the UK Biobank to derive a multimodal structure-function template that captures normative patterns of joint variation in gray matter volume and functional connectivity. We then use this template to perform guided fusion in two independent datasets for evaluation. The resulting ICs show high correspondence with the template, even in modest sample sizes, demonstrating the stability and generalizability of the approach. Finally, we show that subject-specific loadings from template-derived ICs are significantly associated with age, indicating our multimodal normative model captures biologically meaningful variation across the lifespan.

## 2. DATA AND METHODS

### 2.1. Data

We utilize resting state fMRI (rsfMRI) and sMRI data from the UKBiobank (UKB) and Human Connectome Project (HCP) datasets for template generation and validation, respectively. Preprocessed rsfMRI scans (TR = 0.735s; 490 TRs = 6 min total) were downloaded from UKBiobank. Preprocessing details can be found at (https://biobank.ctsu.ox.ac.uk/crystal/crystal/docs/brain_mri.pdf). We applied EPI normalization to MNI space, resliced to 3 x 3 x 3 mm^3^ isotropic voxels, and applied smoothing with a Gaussian kernel with a full width at half maximum (FWHM) of 6mm to the preprocessed data. From HCP, we also obtained preprocessed rsfMRI data for 4 scans (TR = 0.72s; 1200 TRs = 14.4 min total per scan). Preprocessing details for HCP rsfMRI can be found in (http://www.humanconnectomeproject.org/data/).

Preprocessed data were normalized to MNI space, resliced to 3 x 3 x 3 mm^3^ isotropic voxels, and smoothed with a Gaussian kernel (FWHM = 6mm). In both datasets, subjects with head motion larger than 3° rotations or 3 mm translations or with mean framewise displacement larger than 0.3 mm were excluded. For all processed rsfMRI scans, spatially constrained ICA was applied using the GIFT toolbox (https://trendscenter.org/software/gift/) and the NeuroMark 1.0 template [5] to extract subject-level spatial maps and for each of the 53 intrinsic connectivity networks (ICNs) defined in the template, as well as their respective activation time courses. Functional network connectivity (FNC) was computed using pairwise Pearson correlation of time courses for all pairs of ICNs. In HCP, we generated an additional combined FNC feature labeled ‘ALL’ by averaging the FNC matrices from the 4 independent scans.

In UKB, T1-weighted sMRI images were preprocessed with SPM12, and modulated gray matter probabilistic segmentation maps were extracted to obtain gray matter volume (GMV). GMV maps were then smoothed with a Gaussian kernel with FWHM = 12mm. In HCP, T1-weighted sMRI images were preprocessed using the DARTEL tool from SPM12. Image registration, bias correction, and tissue clarification were performed using the unified model. DARTEL was employed to estimate the gray matter volume (GMV) images, which were resliced to a resolution of 1.5 mm and smoothed with a Gaussian kernel with FWHM = 10 mm.

Finally, in UKB we excluded subjects with self-reported psychiatric conditions or symptom of depression and anxiety as described in the minimal phenotyping set defined in (Cai et al., 2021) from the full set of 39,760 QC-passed subjects to derive a final set of 16,530 “unaffected” control subjects. In HCP, we selected a reliable subset of 833 subjects that had data for all 4 scans.

### 2.2. Joint ICA

We utilized the joint ICA (jICA) approach for blind fusion of FNC and GMV maps in UKB to generate the multimodal templates (Fig. 1). Suppose a set of ‘true’ underlying neuronal sources **s**. Given the dataset **x** = [*x*^1^, *x*^2^, …, *x*^*M*^] comprised of (*m* = 1, 2, …, *M*) modalities, where each 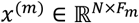 contains *Fm* features from *N* subjects, jICA models the sources **s** as **x** = **As**, where **A** ∈ ℝ^*N*×*C*^ is a shared mixing matrix capturing subject-level loadings across modalities, and each 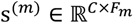 contains modality-specific independent components (ICs). The goal of jICA is to estimate an unmixing matrix **W** (the inverse of **A**) such that **y** given by **y** = **Wx** is a good approximation to the ‘true’ sources **s**. In this work, we employ jICA using the Infomax algorithm [9] with 100 runs, selecting the best run for each.

**Fig. 1.**
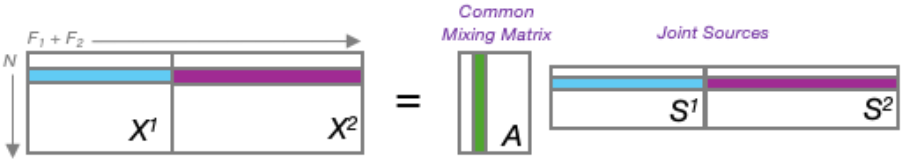
Joint ICA model.

### 2.3. Identifying Reliable Multimodal Templates

We iterated over 5 sample sizes and 11 model orders to identify an optimal set of joint sources that were reproducible across batches. For each sample size in [N = 1000, 2000, 3000, 4000, 5000] we randomly sampled without replacement from the unaffected subjects to generate two non-overlapping batches of size N. We then applied jICA (**2.2**) to each batch at each sample size across model orders [C = 5, 8, 10, 12, 15, 18, 20, 25, 30, 35, 40] for a total of 110 joint fusions. Joint sources were matched across batches for each sample size and model order using a greedy matching approach on the average Pearson correlation of spatial maps for the FNC and GMV components. Correlations were averaged so the joint source correspondence and subsequent matching considered both modalities equally rather than be dominated by the GMV components with much larger dimensionality. We identified the optimal sample size and model order as the set with the highest cross-batch correspondence across all joint sources to serve as the multimodal template.

### 2.4. Multimodal Template Validation

The joint sources from Batch1 of the optimal sample size and model order experiments served as our multimodal template. We performed validation of the template in two data sets: 1) a random sample of equal size from the remaining subjects not used in Batch1 or Batch2 for template generation, and 2) a novel set of N=833 subjects from the HCP dataset.

To apply our multimodal template in new data, we utilized the multivariate objective optimization ICA with reference (MOO-ICAR) approach [5]. MOO-ICAR is a ‘semi-supervised’ constrained ICA approach that utilizes spatial priors (i.e. templates) to guide the decomposition toward a desired set of ICs by solving the dual objective of maximizing both independence of ICs and their correspondence to the template spatial maps. We implemented group-level MOO-ICAR by stacking FNC + GMV inputs across subjects to derive a shared set of joint source IC maps (**s**) and subject-specific loadings (**A**).

In the UKB Batch3 validation experiments, we tested the effect of sample size on the group-level joint source estimation by applying MOO-ICAR on subsets of [N = 100, 250, 500, 750, 1000, 2000, 3000, 4000, 5000] subjects, such that the subjects in the N=100 subset are the same as the first 100 subjects in the N=250 subset, and so on. We compute the Pearson correlation of the joint source maps between each subset and the multimodal template, as well as pairwise across all subsets. We also use linear regression to test the association of the estimated subject loadings with age, controlling for sex.

In the HCP validation experiments, we apply the model on a sample from a novel population of younger subjects than those in the UKB sample used for template generation. We apply group-level MOO-ICAR to estimate sources fusing FNC from each of the 4 scans, as well as the ‘ALL’ FNC averaged across all scans, with the GMV maps. We test the reliability of our multimodal template by computing the correspondence of source maps using repeated scans across the same sample via pairwise Pearson correlation across all scans, as well as correspondence to the template. Again, we use linear regression to test the association of the estimated subject loadings with age, controlling for sex.

## 3. RESULTS

We computed the cross-batch correspondence of each matched pair of ICs across our 110 fusion experiments corresponding to varying sample size and model order (Fig. 2). We observed increasing cross-batch IC correspondence with increasing sample size. While smaller model orders (<10) had high correspondence across all ICs in the 4K and 5K sample sizes, visual inspection showed reduced granularity in the GMV ICs specifically, so we focused the search on the larger model orders (>20). We found the 5K sample and model order = 30 components yielded the highest cross-batch correspondence (*r* = 0.64-0.94, *r*mean = 0.83) was selected as the multimodal template (Fig. 3).

**Fig. 2.**
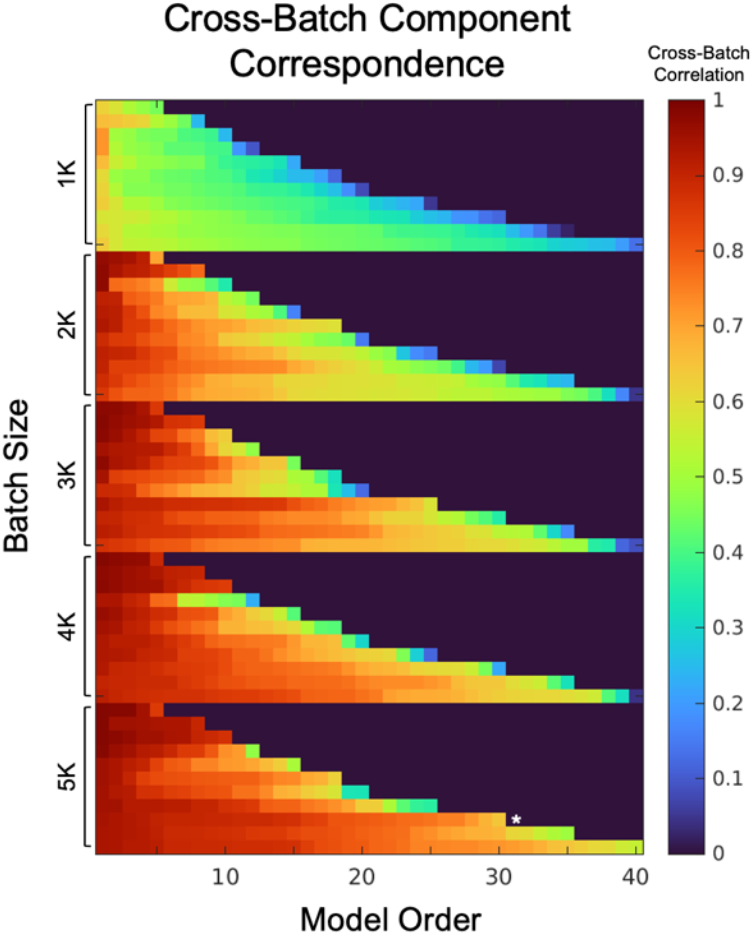
Cross-batch correspondence of IC maps identified batch size = 5000 subjects, model order = 30 components as the optimal multimodal template (denoted with *).

**Fig. 3.**
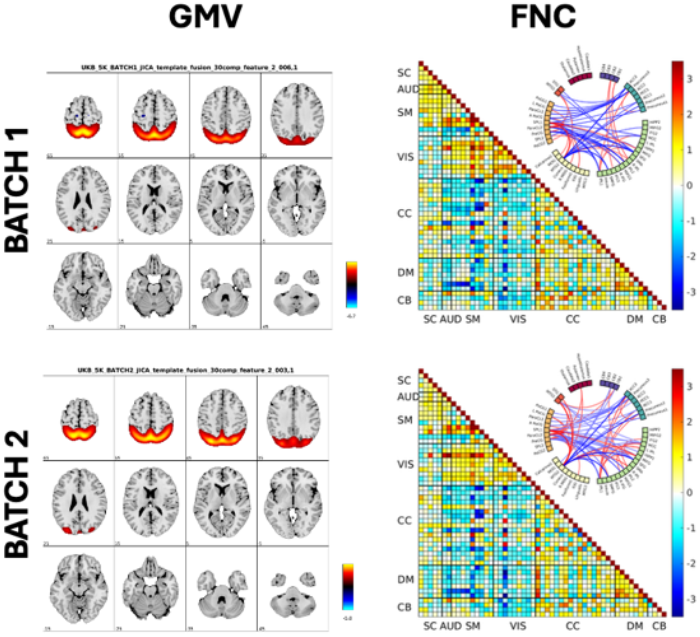
Top matched ICs between Batch1 & Batch2 (*r* = 0.94).

Validation in UKB Batch3 shows strong correspondence of MOO-ICAR estimated ICs with the template and across subsets even at small sample sizes, with fair correspondence (*r* = 0.78) at N = 100 subjects and excellent correspondence (*r* > 0.91) at N >= 250 subjects (Fig. 4A). When viewed at the individual IC level, a majority of components at N=100 subset showed fair to good (*r* = 0.70–0.90) correspondence with template spatial maps, with only the last component having relatively poor correspondence (*r* = 0.45). All other components at N>=250 showed good to excellent correspondence (*r* > 0.69) (Fig. 4B). Regression analyses found 29/30 ICs (all but IC18) exhibited significant (pFDR < 0.05) associations with age (adjusted for sex) (Fig. 6).

**Fig. 4.**
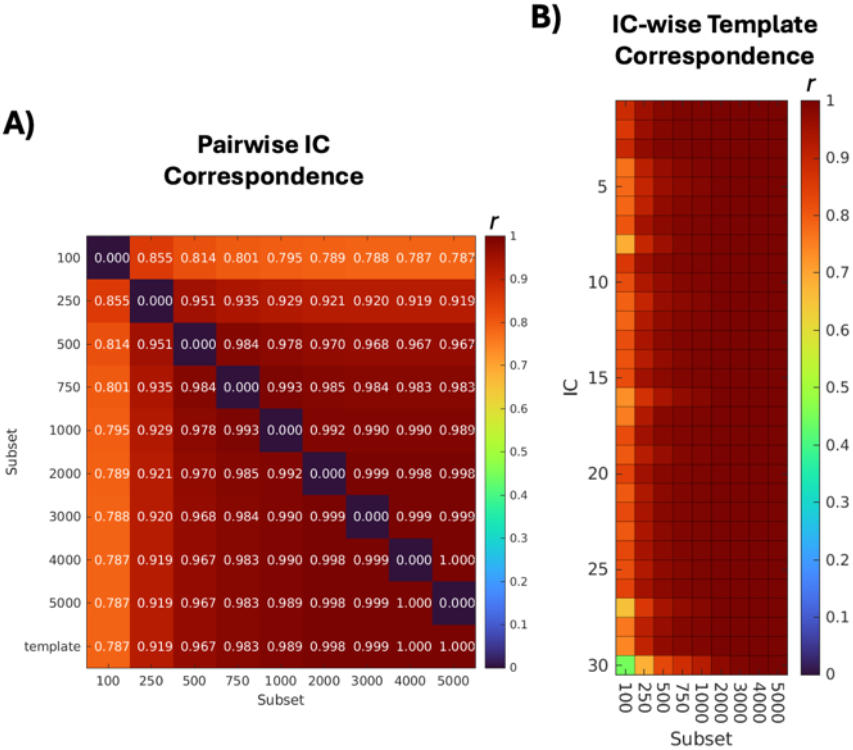
UKB results with A) pairwise correspondence of joint source maps across subset sizes and B) IC-wise correspondence of joint source maps to template ICs. Results show high correspondence with sample sizes >100 subjects.

Validation in HCP showed near-perfect pairwise correspondence (r > 0.99) of group level component maps across scan experiments, and good correspondence (r > 0.81) between all scan experiments and the multimodal template (Fig. 5A). IC-wise correspondence showed good (r = 0.64-0.89) correspondence to template that was highly stable across scan experiments (Fig 5B). Associations of subject-level loadings with age (controlled for sex) were also stable across scan experiments, with 14/30 ICs exhibiting significant (pFDR < 0.05) relationships (Fig. 6).

**Fig. 5.**
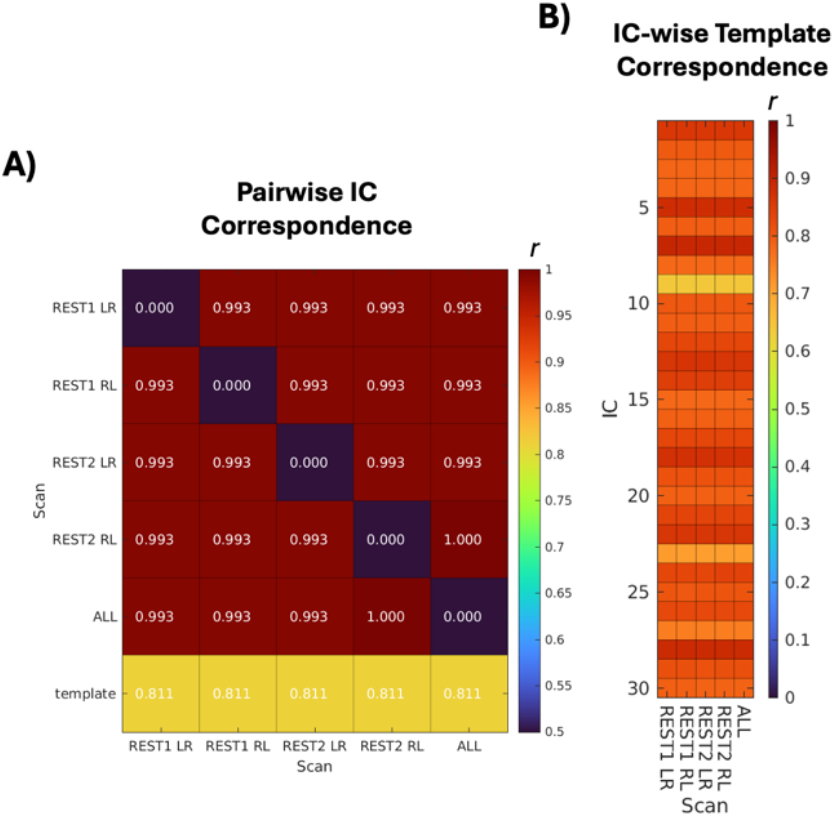
HCP results with A) pairwise correspondence of joint source maps across scans and B) IC-wise correspondence of joint source maps to template ICs. Results show high correspondence across scans and good correspondence to template in new population.

**Fig. 6.**
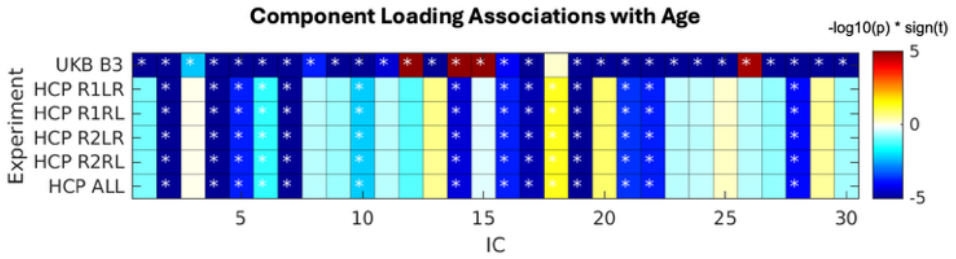
Associations (-log10(pfdr) * sign(t)) of subject-level IC loadings with age (controlled for sex) for UBK Batch3 and HCP experiments. (*) denotes significance (pfdr < 0.05).

## 4. DISCUSSION

We present a multimodal FNC + GMV normative template for reliable and replicable structure-function fusion. In our template generation experiments, we found relatively low cross-batch correspondence in a majority of ICs between independent fusions when sample sizes were “small” (i.e., N = 1000), even when batches were both derived from the same UKB dataset (Fig. 2). Given that samples of this size are already considered large in most neuroimaging studies, these results motivate the need for a standardized multimodal normative template that can be applied consistently across studies. However, a few ICs have high cross-batch coherence at this batch size, which corresponds to previous work in which we showed ICs in certain brain regions are highly stable, i.e. less influenced by linkage between structure and function, whereas a majority of ICs have spatial maps that are strongly influenced by cross-modal relationships [10]. Joint ICs become more generalizable as sample size increases, indicated by higher cross-batch correspondence (Figs 2-3).

Validation experiments in UKB Batch3 shows high correlation to the template at truly small samples (N = 100), and excellent coherence to the template when N >= 250 subjects are used, further illustrating the utility of our multimodal template at small samples (Fig. 4). HCP validation experiments also show good correspondence (r = 0.81) to the template at a moderate sample size (N = 833), as well as excellent coherence (r > 0.99) within the sample across scans (Fig. 5). Finally, significant associations between subject-level loadings and age in all validation experiments indicate the template-derived ICs capture biologically relevant variation in multiple populations (Fig. 6). Future work will extend this approach to clinical populations to evaluate whether individual deviations from the multimodal normative model capture cognitive, behavioral, and symptom-level variation across psychiatric diagnoses. Methodological extensions may also focus on expanding the MOO-ICAR framework to estimate subject-specific multimodal spatial maps, enabling more granular normative deviation mapping beyond group-level representations.

